# Nanopatterned Thermoresponsive Functionalization of Substrates via Nanosphere Lithography

**DOI:** 10.1101/796268

**Authors:** Marcela Mireles, Cody W. Soule, Luis F. Delgadillo, Thomas R. Gaborski

## Abstract

Self-assembled monolayers (SAMs) have been widely utilized as a way of tailoring surface chemistry through the adsorption of organic molecules to different materials. SAMs are easy to prepare and offer a wide variety of organic molecules that afford additional or improved properties to the coated material. Spatial control of SAM placement has been achieved over many length-scales, even at the nanoscale. However, nanopatterned SAMs are usually prepared through serial processes utilizing atomic scanning probes or soft-lithography utilizing elastomeric masters. These techniques are expensive or not repeatable. Here we present the use of nanospheres for the creation of nanopatterned Au:Cu films which spatially control the grafting of a thermoresponsive SAM made from poly(N-isopropyl acrylamide) (PNIPAM). Chemical characterization validates the presence of PNIPAM and environmental atomic force microscopy showed its response to temperature which was evidenced by a change in stiffness. Our approach represents an affordable large area methodology for repeatable spatial control of SAMs at the nanoscale.

## 2 Introduction

Surface chemistry engineering is the essential task of enhancing or improving the properties of materials through the manipulation of its properties at the surface. Thin film coatings of inorganic and organic materials have been commonly used to tailor surface properties. These films are deposited onto the desired surface by varied physical and chemical methods. Among these approaches, self-assembled monolayers (SAMs) are of particular interest due to ease of processing amenable to atmospheric conditions. SAMs consist of molecules that spontaneously organize over a substrate as a tightly packed monolayer.^1^ The molecules display a head group with high affinity for the substrate which, acting as an anchor, is responsible for the substrate-molecule interaction determining the molecular orientation. The head group is either attached to a functional group through a spacer or is directly linked to a functional molecule. This molecular anatomy offers tremendous versatility as a wide variety of functional groups and molecules can be utilized.^2,3^

The ability to control the spatial arrangement of SAMs along the plane of the monolayer has been sought throughout the last three decades.^1,4,5^ The routes for the patterning of SAMs can be divided into those that manipulate the SAM molecules themselves (direct) and those that manipulate the surface presented to the SAM molecules (indirect). Direct approaches are those that directly place or replace SAM molecules over the substrate and can be further divided into serial and parallel processes. Scanning probe-based methods utilize scanning probe or e-beam microscopes offering excellent control and achieving nanoscale resolution.^6–8^ However, these are time consuming serial processes that significantly increase fabrication costs and therefore are not used for practical applications requiring large areas. Soft lithography-based methods utilize elastomeric masters as a way to deliver the SAM molecules over a spatially controlled pattern.^9,10^ Although this is a parallel process that can achieve nanoscale resolution, its application to large scale remains halted due to fundamental and practical issues.^11,12^ Furthermore, the nanoscale masters utilized are fabricated through serial processes themselves. Indirect approaches are those that modify the substrate by presenting regions of high and null affinity towards the SAM molecules and can be further divided into serial and parallel processes. An example of this affinity is the sulfur-metal chemistry which has been the subject of intense research and is well understood and applied.^13–16^ Thiol terminated molecules show high affinity for transition metals over oxides. Creating a two dimensional metallic pattern over a surface and exposing to thiol-terminated molecules can produce a spatially controlled SAM. E-beam lithography represents the main serial process used to pattern a surface. Compared to parallel process offering control and nanoscale resolution, e-beam lithography is time consuming and expensive. In contrast, traditional photolithopgraphy is a parallel inexpensive process, however does not offer nanoscale resolution.

The ability to pattern SAMs with nanoscale resolution over a large area through a fully parallel process remains a challenge. The use of nanospheres as pattering features, nanosphere lithography (NSL), is a bottom-up approach that takes advantage of the self-organization of nanospheres at varied interfaces, offering reliability and repeatability over large areas.^17–19^ Here we present the use of NSL as way of creating a metallic nanoscale pattern over a substrate which can then be spatially functionalized with thiol-terminated functional molecules. A similar approach has been previously published using a nanopattern of Au over or under TiO_2_ and functionalized with an alkanethiol.^20^ In the present work the metallic layer was a thin film of Au:Cu, which a showed higher grafting ratio than Au only (**Figure S1 – Supporting information**). The Au:Cu nanopattern presented here can be utilized to spatially pattern SAMs with many different functional groups as long as the thiol headgroup remains present. We chose to graft a thiol-terminated poly(N-isopropyl acrylamide) (PNIPAM) as this is a smart thermoresponsive polymer exhibiting a conformational globule/coil transition. When grafted onto a surface, the PNIPAM molecules self-assemble as an organized brush. Above 32 °C, the brush consists of collapsed PNIPAM molecules (globule) which unfold and extended (coil) as the temperature moves below 32 °C; this temperature is known as the lower critical solution temperature (LCST).^21–23^

Uniform PNIPAM coatings have been used to cell adhesion/detachment^24–28^ and for self-cleaning antifouling coatings of sensors and membranes.^29–33^ Nanopatterned PNIPAM has the potential of controlling cellular adhesion at the nanoscale improving cell sheet characteristics and potentially eliciting alignment. Moreover, and added benefit of the grafted PNIPAM is the ability of adjusting the effective stiffness sensed by the cells. Stiffness has been found crucial in stem cell differentiation, cell sheet harvesting substrates with adjusted effective stiffness are ideal for tissue engineering. In the case of sensors a dense but non-continuous PNIPAM coating would display self-cleaning capabilities without compromising sensing area. Furthermore, nanosphere lithography has also been utilized for the fabrication of nanoporous membranes by our group.^34^ The technique presented here serves as an extension enabling the functionalization specifically of the surface and not the pore walls. This advantage may be of particular value to large area nanoporous membranes with precisely engineered pore properties that would otherwise be altered and compromised by a conformal polymer coating.^33,35–40^

Our aim is to provide a straightforward methodology for the indirect patterning of SAMs at the nanoscale. The approach we present can be used for a wide variety of SAMs. Here the metal-S affinity was employed but the method is not exclusive to this chemistry and can be applied to other compatible SAM-substrate systems. Presence of PNIPAM on the functionalized samples was demonstrated physically by a decrease in roughness and chemically by chemical surface analysis. Moreover, the conformational transition was observed when imaging its surface in an environmental atomic force microscope with which stiffness change was detected upon temperature variation.

## 3 Experimental section

We chose a bottom-up fabrication approach to achieve selective functionalization. We used a thiol headgroup than can spontaneously absorb to clean metals and form a SAM.^13^ Figure 1 shows the overall process followed to generate the Au:Cu nanopattern used to indirectly control the spatial arrangement of the SAM used. In the final substrate the thermoresponsive polymer exhibits empty circular areas (holes or pores). The inverted pattern (pillars or posts) can be achieved by using the nanospheres to create an etching mask over the Au:Cu. Our group has previously reported on the use of nanosphere lithography for the fabrication of nanoporous membranes,^34^ the process depicted in Figure 1 has been adapted to yield nanopatterned SAMs.

**Figure 1.**
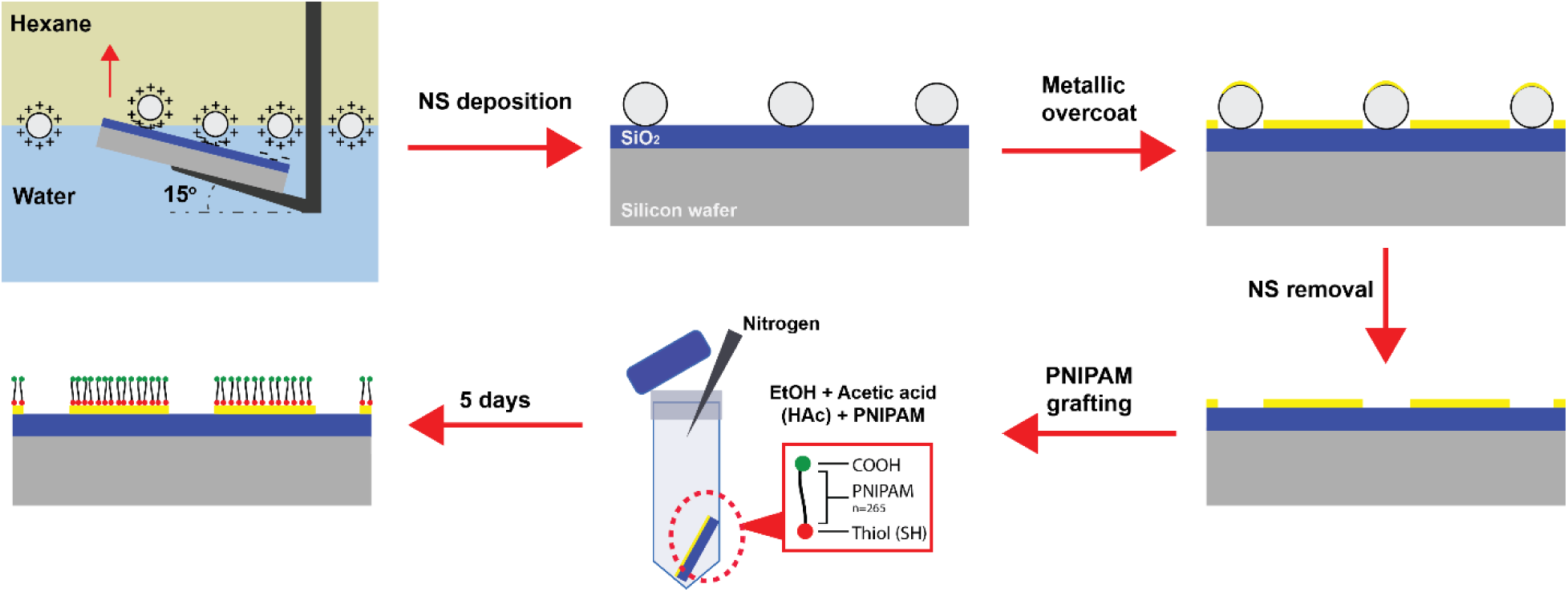
Spatially controlled nanopatterning of PNIPAM. Self-assembled nanospheres are deposited on the substrate and used as patterning features for the Au:Cu metallic overcoat. The thiol-terminated PNIPAM preferentially attaches to the Au:Cu and not to the silicon oxide substrate.

### 3.1 Fabrication of the Au:Cu Nanpatterned Substrates

Positively charged 300 nm polystyrene nanospheres were allowed to self-assemble at a hexane/water interface through which the substrate was pulled across at speed of 250 μm/s resulting in deposition of a monolayer of nanospheres.^17^ The substrates used consist of a silicon wafer coated with silicon oxide deposited by plasma enhanced chemical vapor deposition to a thickness of 300 nm and annealed at 600 °C for 1 hr under nitrogen flow.^41^ Next, a thin layer of Cr (3 nm) is deposited by e-beam evaporation as an adhesion layer for the Au:Cu. Then, 13 nm of Au:Cu were deposited by sputtering at 200 W (DC) in an Ar atmosphere with a working pressure of 3 mTorr. The elemental composition of this layer as analyzed by X-ray photoelectron spectroscopy (XPS) and has been included in the **supporting information (Figure S2)**. Finally, the nanospheres were removed in toluene under sonication for a total of 16 mins and cleaned sequentially in acetone, isopropanol, and ultrapure water for 4 mins each under sonication. The metallic deposition was also done over SiO_2_ substrates without nanospheres (plain substrates). All substrates, nanopatterned and plain, were annealed at 250 °C for 3 hrs over a hot plate inside an enclosed fume hood at atmospheric conditions.

### 3.2 PNIPAM Grafting

PNIPAM with a molecular weight of 30,000 g/mol exhibiting a thiol and a carboxylic acid group on either end was used as purchased from Polymer Source (Québec, Canada). The grafting process, adapted from the literature, undergoes in ethanol to which acetic acid (HAc) was added to avoid the formation of multi-layer structures by passivating the carboxylic acid groups in the α-carboxy ω-thiol terminated PNIPAM used.^42,43^ The samples were submerged in a 100 µM solution of PNIPAM in ethanol with a varying fraction of HAc (0, 2, and 10% by volume) for 5 days. The extended self-assembly time promotes a densely packed, well ordered monolayer, while the acetic acid in the solution bonds to the carboxyl end groups on the PNIPAM chain, ensuring that formation of a multilayer structure will not occur.^44^ The grafting is performed in a 50 ml centrifuge tube filled with nitrogen, sealed with parafilm and kept in a dark environment. After self-assembly, the excess polymer chains are removed by multiple alternating rinses in ethanol and ultrapure water.

### 3.3 Chemical Characterization

PNIPAM functionalized plain substrates were used to study the chemical composition at the surface by XPS using a Kratos AXIS Ultra DLD system. The tool is equipped with an Al-Kα source and the take-off angle was set to 70° in order to enhance surface signal. The data obtained was analyzed with the CasaXPS processing software.

### 3.4 Surface Characterization

#### Plain substrates

The topography of theplain substrates was recorded by atomic force microscopy (AFM) using an equipment from NT-MDT Spectrum Instruments (Moscow, Russia). The tool is equipped with a triangularly shaped Si_3_N_4_ cantilever used in non-contact (tapping) mode. The data obtained was analyzed with the open source Gwyddion software.^45^

#### Nanopatterned substrates

The topography of nanopatterned substrates was recorded on an environmental AFM (MFP-3D) from Oxford Instruments NanoAnalysis & Asylum Research (Santa Barbara, United States). The samples were affixed to a fluid cell filled with DIW, the cell is equipped with a heating element allowing precise control of the temperature. Measurements were taken at 23 °C (RT) and 37 °C, after incubating the samples for 2 hrs to allow the PNIPAM to collapse or extend.

### 3.5 Use of Polystyrene Beads to Detect PNIPAM Globule/Coil Transition

Plain samples with and without PNIPAM were placed inside a device which had a top channel in contact with the surface of the sample. The device was set on a hot plate to be maintained 37 °C temperature and DI water was flowed for 30 mins to prime the surface. Then, a solution containing 1 μm fluorescent polystyrene beads was flowed at 10 μl/min for 10 mins. Next, the samples were rinsed with DI water to remove floating beads, then air was introduced and the sample was allowed to dry on the hot plate. The sample was then imaged in a fluorescent microscope (Leica Microsystems, Wetzlar, Germany) to record the number of beads non-specifically attached to the sample. Next, the sample was placed in DI water at RT (23 °C) for 30 mins to allow the PNIPAM to extend and release attached beads. The sample was then imaged to record the number of remaining beads.

### 3.6 Young’s Modulus Evaluation

Using the same environmental AFM tool described above the Young’s modulus was evaluated at 23 °C and 37 °C. The samples were allowed to reach thermal equilibrium at the desired temperature for at least 2 hr. Then, a 20×20 μm scan was recorded to select a non-porous area of 2×2 uμ on average from which 100 force curves were recorded arranged on 10×10 matrix. Asylum Research software was used to fit the raw data and obtain a histogram of Young modulus values for each sample measured.

## 4 Results and Discussion

First, we utilized plain Au:Cu samples to evaluate the presence of HAc in the grafting solution by recording the topography and chemical composition at the surface. We then monitored the interaction of polystyrene beads with the grafted PNIPAM at different temperatures. Lastly, we utilized nanopatterned samples to study the effective stiffness of the PNIPAM in its globule and coil states. In this section we present and discuss the results obtained.

Oxidation of the Cu in the Au:Cu film is expected to happen when in contact with the environment and HAc chemically etches oxidized copper. We first needed to optimize the HAc concentration to allow for the removal of oxidized copper at the surface, but also to prevent an excess of HAc from penetrating further into the layer which could potentially cause delamination. We evaluated the concentration of HAc by adding 0, 2, or 10% by volume to the grafting solutions. Plain substrates were functionalized and characterized by AFM and XPS. It has been shown before that a decrease in roughness, from the metallic to the functionalized surface, can be correlated with the presence of a SAM.^30,46^ Roughness decrease is a result of uniform and conformal coverage with macromolecules that effectively shadow or planarize the substrate. Our results are shown in Figure 2 with the largest reduction, from 5.0 to 3.3 nm, obtained for the 2% HAc sample, which suggests this to be the optimal concentration for conformal coverage. On the contrary, the sample functionalized without HAc exhibits similar roughness to the bare Au:Cu sample, suggesting that if any grafting took place it resulted in a non-uniform or sparse coverage.

**Figure 2.**
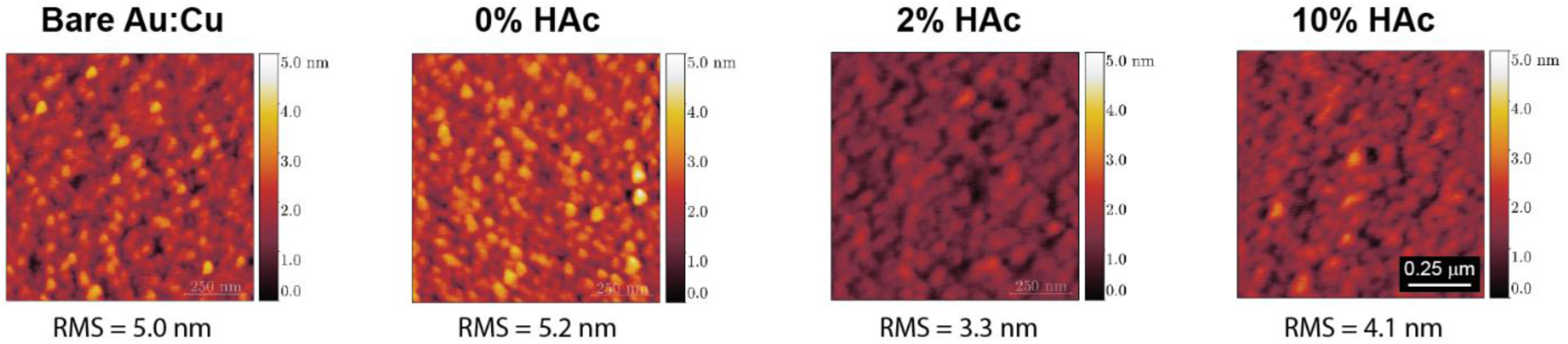
2D topography scans. Bare Au:Cu sample and PNIPAM functionalized with 0, 2, and 10% HAc. The roughness has been annotated at the bottom of each scan and expressed as RMS.

Next, the samples were analyzed by XPS to characterize their chemical composition. The samples were not sputter cleaned prior to recording the spectra and were measured as introduced into the XPS chamber. Figure 3 shows the XPS data obtained from the Cu_2p_ and S_2p_ regions for a plain Au:Cu substrate as well as functionalized plain substrates for the HAc concentrations tested. The plain Au:Cu and the 0% HAc samples show the presence of Cu-O (Cu_2p3/2_ = 933.6 eV)^47,48^ which is expected since oxidation of Cu readily happens in air. However, the samples functionalized in the presence of HAc show virtually no Cu-O, this is due to the fact that HAc etches oxidized Cu at the surface exposing metallic Cu that can interact with the thiol-terminated PNIPAM. All of the functionalized samples show presence of Cu-S (Cu_2p3/2_ = 932.0 eV)^47,49^ which indicates grafting has taken place. The 2% HAc sample showed the largest Cu-S peak which is indicative of a denser SAM on this sample and is in good agreement with the data found for roughness. The data obtained from the S_2p_ region is also shown in Figure 3 where absence of a peak at high binding energy (163 eV) is indicative of successful prevention of a multilayer coating by the HAc passivating the carboxyl group on the PNIPAM.^42,44,50^ The S_2p_ data was deconvoluted into two main components centered at 161.1 and 162.3 eV. It has been well established that metallic thiolate bonds,^14,42,44,50,51^ such as Au-S and Cu-S, result in a S_2p3/2_ peak located at 162 eV which is in good agreement with our data. The peak observed at a lower binding energy has been associated with degradation during XPS due to cleavage of the C-S bond.^51^ Degradation processes caused from irradiation during XPS analysis result in the formation of new peaks with increasingly larger height after each scan. This change in the spectra, that can be readily identified, was not observed during the collection of the data presented in Figure 3. This peak has also been attributed to the presence of additional sulfides where more than one thiol head interact with the Au surface.^14,48,49,51,52^ The chemistry at the interface of thiol-terminated SAMs and metallic substrates remains a matter of debate and often requires more sophisticated characterization tools than XPS.^53^ Nevertheless, the data presented here is sufficient for the purpose of establishing the twofold role of HAc and the presence of a chemisorbed SAM. The overall area under the curve for the 0% HAc sample indicates this to be the least dense film which is in agreement with the lack of roughness decrease found for this sample.

**Figure 3.**
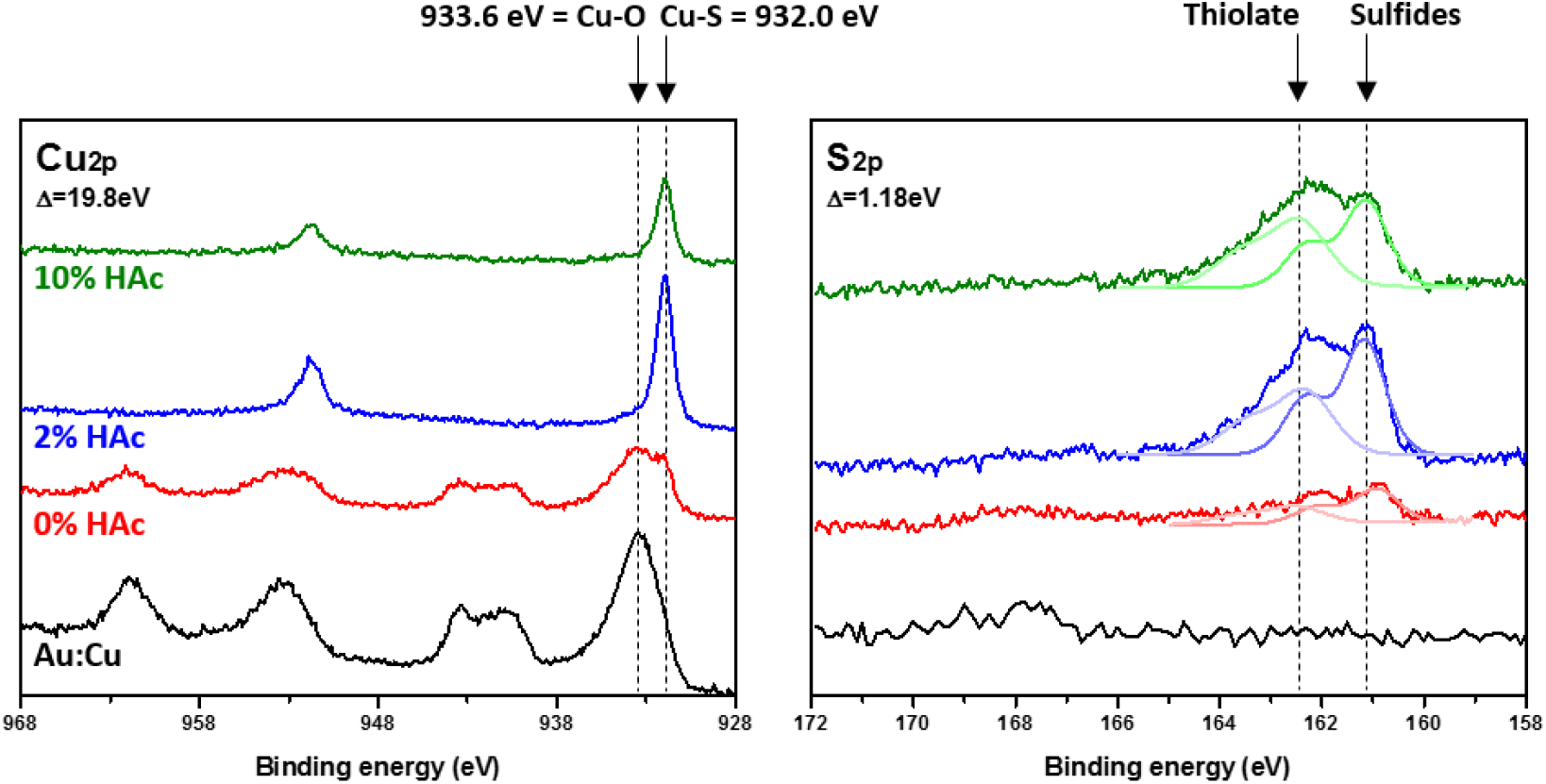
Chemical characterization. XPS results obtained for PNIPAM functionalized Au:Cu plain substrates with 0, 2, and 10% HAc as well as non-functionalized. The regions corresponding to Cu_2p_ and S_2p_ are presented with the main chemical identities labeled.

To evaluate the presence of the PNIPAM and its functionality we designed a simple test described as follows. Non-functionalized (labelled as bare Au:Cu) and functionalized plain substrates with 2% HAc (labelled as PNIPAM) were placed in a silicone device with a top channel through which a solution containing fluorescent polystyrene beads was flowed for 30 mins at 37 °C. Next, the devices were cooled down to RT and rinsed with DI water. Figure 4b shows the number of beads found on the samples tested. At 37 °C fewer beads are able to attach to the PNIPAM sample, this is in agreement with the antifouling property of this material. A decrease in the number of attached beads was recorderd upon cooling the devices to RT. The difference expressed as a percentage has been plotted in Figure 4c which shows that for the PNIPAM sample, nearly 80% of the beads dettached from the surface as the temperature of the device shifted to RT. This dettachment suggests that as the PNIPAM transitions from a globule into a coil and extends, the beads are essentially pushed off the surface confirming the presence and functionality of the PNIPAM.

**Figure 4.**
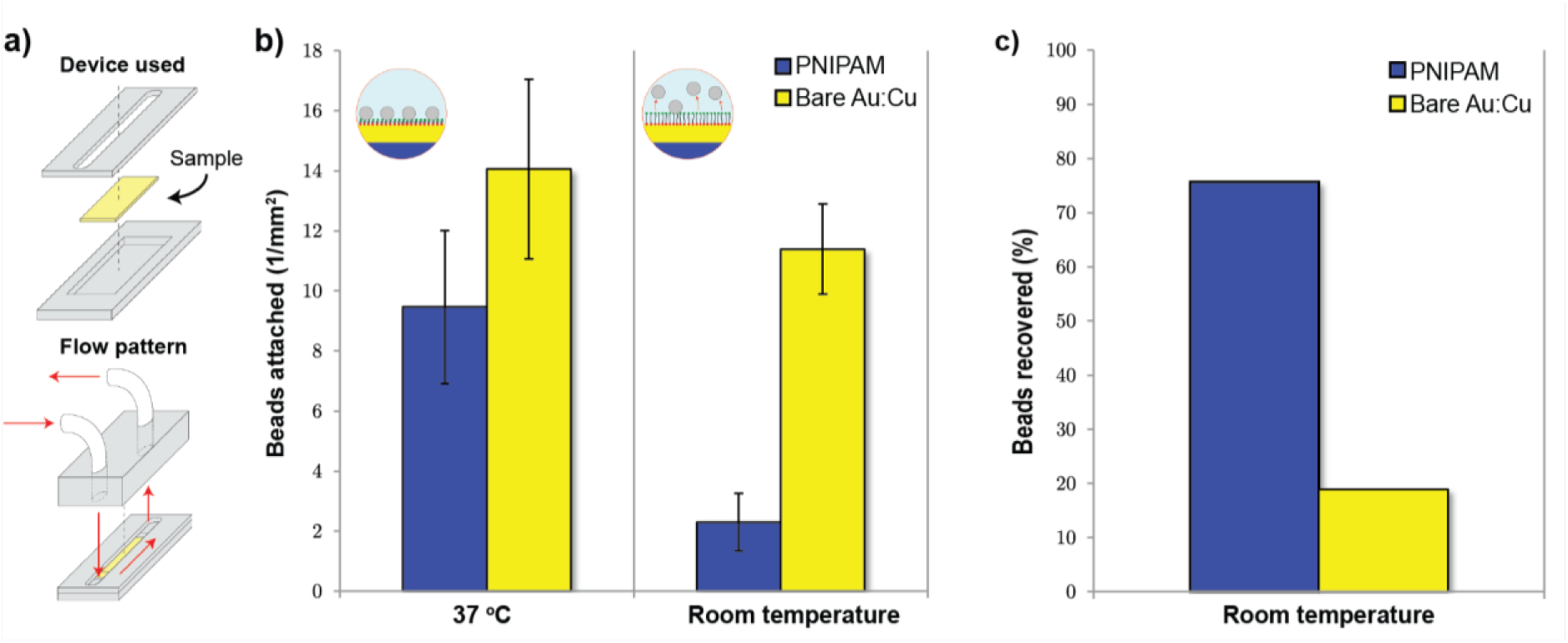
Use of polystyrene beads to detect PNIPAM globule/coil transition. a) device used and corresponding flow pattern, b) density of beads on the surface, and c) percentage of beads detached upon PNIPAM conformational transition.

To further confirm the presence and functionality of the PNIPAM we imaged the topography of the functionalized nanopatterned samples in an environmental AFM from Oxford Instruments (Asylum Research). The AFM is equipped with a closed fluid cell integrated with a heating element allowing for the samples to be imaged while immersed in DI water at RT and 37 °C. The resulting scans are shown in Figure 5 for the two nanopatterned samples tested corresponding to 2 and 10% HAc. The features in the scan appear blurry at RT but sharpen upon temperature increase to 37 °C. We suggest that the blurriness observed is an imaging artifact caused by the AFM tip partially compressing as well as bending the extended PNIPAM at RT. This topographical difference further supports the successful grafting of the PNIPAM onto the nanopatterned surfaces and demonstrates its thermal response.

**Figure 5.**
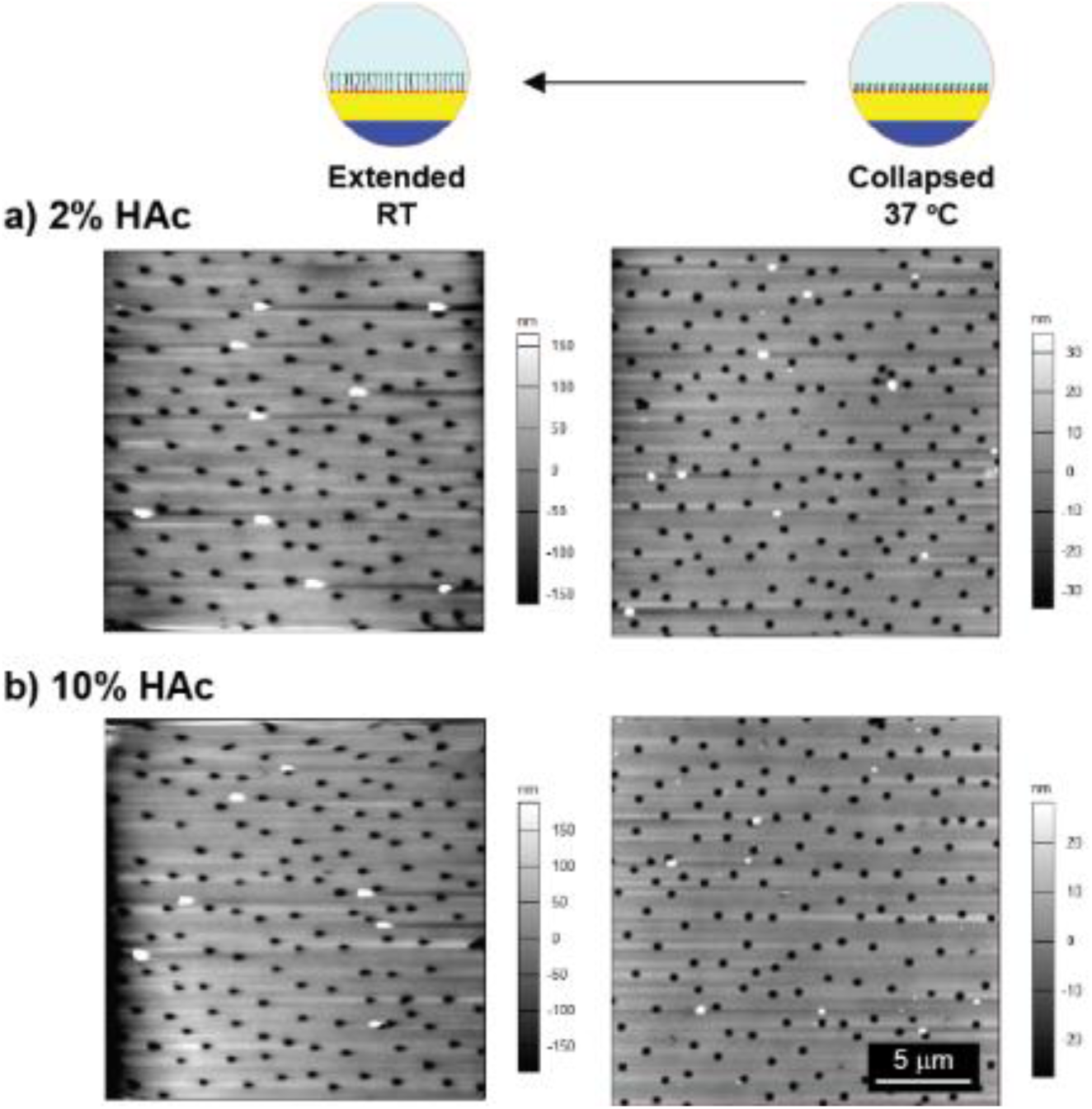
2D topography scans taken in an environmental AFM. Non-contact mode AFM scans were taken in a closed fluid cell chamber at 37 °C and RT for Au:Cu nanopatterned substrates functionalized in a PNIPAM solution containing a) 2% HAc or b) 10% HAc.

To further confirm the successful extension and collapse of PNIPAM on the surface, we measured stiffness changes as a function of temperature. We collected a series of force-indentation curves from a selected area in between the pores and calculated the Young’s modulus using Hertz theory. The obtained values have been plotted as frequency histograms in Figure 6a. As the temperature changes from RT to 37 °C and the PNIPAM transitions from extended to collapsed, the stiffness of the surface is dominated by the PNIPAM or by the underlying substrate, respectively. This behavior was seen for the functionalized samples which showed a higher Young’s modulus at 37 °C with a value similar to that reported for SiO_2_ (10^10^ Pa).^54^ In turn, at RT, both samples displayed a softer surface, however, the 2% HAc registered a larger decrease from 10^9^ to 10^5^ Pa. The larger Young’s modulus shift suggests the presence of a more uniform layer. This difference is consistent with previous results shown in this work indicating the 2% HAc prepared sample has a denser PNIPAM layer. Furthermore, the indentation data (Figure 6b) for this sample at RT shows a gradual change in slope as the tip approaches and retracts, which in conjunction with the hysteresis recorded, indicates a plastic behavior resulting from the tip compressing the PNIPAM extended SAM.^55,56^ Similar behavior in stiffness change with temperature has been reported for PNIPAM nanopillar surfaces.^57^

**Figure 6.**
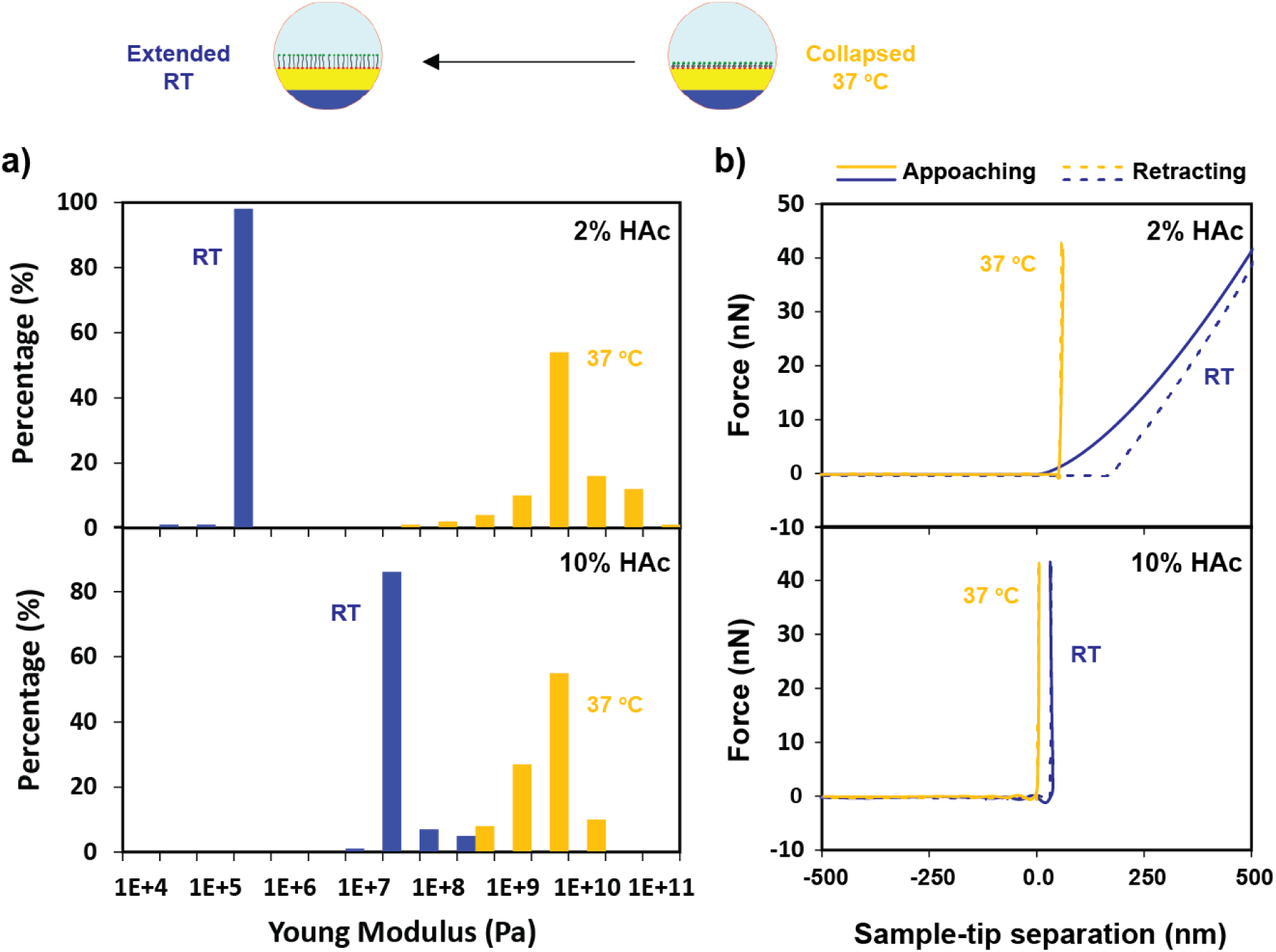
Stiffness evaluation in an environmental AFM. Change in stiffness as calculated from force curves performed in a close fluid cell AFM chamber at 37 °C and RT for Au:Cu nanopatterned substrates functionalized with 2 and 10% HAc. B) Representative force-indentation curves for both 2% and 10% HAc samples at the two different temperatures, tip approach (solid line) and retraction (dotted line) are shown.

## 5 Conclusions

In this manuscript, we presented a methodology for spatially controlling SAMs at the nanoscale. Utilizing nanosphere lithography we created a Au:Cu nanopattern to which the thermoresponsive PNIPAM was grafted. The use of acetic acid in the grafting solution was shown to critical the process by ensuring the formation of a monolayer but more importantly by resurfacing metallic Cu due to *in-situ* copper oxide removal. The latter is essential for the successful implementation of Cu for SAMs. The responsiveness of the polymer as a function of temperature was correlated to stiffness decrease upon polymer unfolding. This stiffness change is due to the compliant nature of the extended PNIPAM. We expect this affordable spatial control of SAMs at the nanoscale to advance the overall field of surface engineering, particularly in applications such as tissue engineering and membrane science.

## Acknowledgements

Research reported in this publication was supported in part by NIGMS of the National Institutes of Health under award no. R35GM119623 to TRG.

