## Supporting information for "Nanopatterned Thermoresponsive Functionalization of Substrates via Nanosphere Lithography"

**Figure S1 Sulfur composition.** Deconvolution of the $S_{2p}$ region for functionalized plain Au only and Au:Cu substrates PNIPAM functionalized with 0, 2, and 10% acetic acid. Overall the signal from Au only samples was lower than from the Au:Cu samples. Moreover, presence of oxide was detected for the Au only sample at 10% HAc.

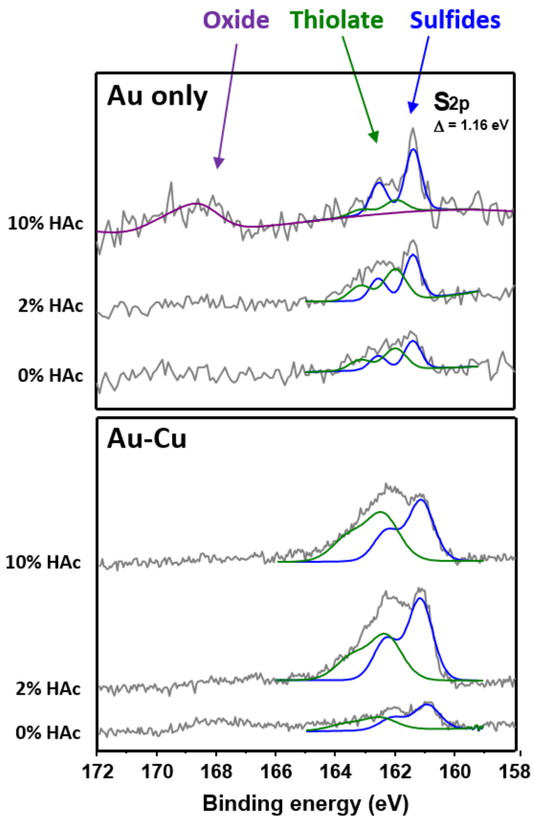

**Figure S2 Au:Cu substrate elemental composition.** As calculated from the XPS regions for Au4f, O1s, C1s, and Cu2p taking into account their respective sensitivity factors. The composition has been expressed in percentage and annotated within each plot.

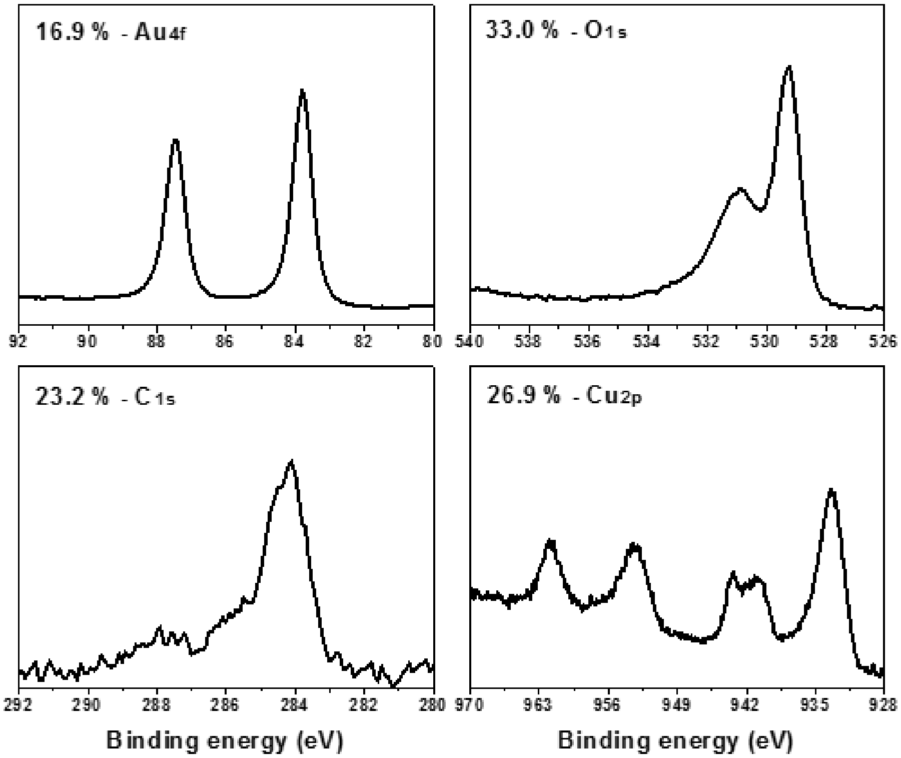
